## Supplemental Information for "Potent AMA1-specific human monoclonal antibody against P. vivax Pre-erythrocytic and Blood Stages"

^1^ Center for Global Health and Diseases, Department of Pathology, Case Western Reserve University School of Medicine, ^2^ Walter and Eliza Hall Institute of Medical Research, Parkville, Victoria, Australia, ^3^ Department of Medical Biology, The University of Melbourne, Parkville, Victoria, Australia, ^4^ Burnet Institute, Melbourne, Victoria, Australia, ^5^ Department of Medicine, The University of Melbourne, Parkville, Victoria, Australia, ^6^ Mahidol Vivax Research Unit, Faculty of Tropical Medicine, Mahidol University, Bangkok, Thailand, ^7^ Department of Microbiology, Monash University, Clayton, Victoria, Australia, ^8^ Malaria Research Unit, Institut Pasteur du Cambodge, Phnom Penh, Cambodia, ^9^ Department of Infectious Diseases, The University of Melbourne, Parkville, Victoria, Australia, ^10^ Central Clinical School and Department of Microbiology, Monash University, Clayton, Victoria, Australia,^.11^ Vaccine and Gene Therapy Institute, Oregon Health & Science University, Beaverton, OR, ^12^ Center for Global Infectious Disease Research, Seattle Children’s Research Insititute, Seattle, WA, ^13^ Department of Pediatrics, University of Washington, Seattle, WA, ^14^ InterRayBio LLC,^15^ Veterans Affairs Medical Center, Cleveland, OH

**Index Supplemental**

**Figures:**

Supplemental Figure 1 – Blocking activity of serum from Cambodian individuals

Supplemental Figure 2 – Flow diagram depicting PvAMA1-specific B cells that were isolated

Supplemental Figure 3 – HumAb Single Cycle Kinetics

Supplemental Figure 4 - Replicates of Pf-PvAMA1 cell line inhibition

Supplemental Figure 5 – Sporozoite HC04 invasion separated CSP210/247

Supplemental Figure 6 – Microscopic assessment of P. vivax liver in FRGN huHep mice after administration of anti-AMA1 human monoclonal antibody 826827

Supplemental Figure 7 - Surface properties of PvAMA1 and its interaction partners RON2 and humAb 826827

Supplemental Figure 8 – RON2 binding groove conservation and Sequence differences between Pv, PvPNG16, Pf, Pc, Pk, Tg

Supplemental Figure 9 – Published PvAMA1 Clinical Isolates sequence conservation

Supplemental Figure 10 - Epitope and paratope of the PvAMA1-826827 complex

**Tables:**

Supplemental Table 1 – VDJ alleles and CDR3 sequences and clonal groups

Supplemental Table 2 – Avidity Index (AI_50_) of PvAMA1 specific humAbs to different AMA1 constructs

Supplemental Table 3 - List of SNP and haplotypes observed in AMA1 sequences in isolates tested in response to 826827

Supplemental Table 4 – Crystallography data collection and refinement statistics

Supplemental Table 5 - Interactions PvAMA1 – Fab 826827 based on PISA

**Movies:**

Supplemental Movie 1: Morph between open and closed conformation of AMA1 Domain Loop 2

Supplemental Movie 2: PvAMA1 bound Fab826827 compared to RON2

Supplemental Movie 3: Polymorphisms within PvAMA1 in relation to Fab826827

**Supplemental Tables & Figures**

**Supplemental Figure 1 – Blocking activity of serum from Cambodian individuals**

Inhibition of PvRON2 binding to PvAMA1 by plasma samples from Cambodian donors. Plasma samples were tested at 1/50 dilution for the ability to inhibit binding of the PvRON2 loop to PvAMA1 in a plate-based assay. Samples were tested in duplicate. Data show mean and range; the dotted line shows 50% inhibition. PBMCs from donor C5 were selected for sorting B cells specific for PvAMA1 and subsequent MAb generation.

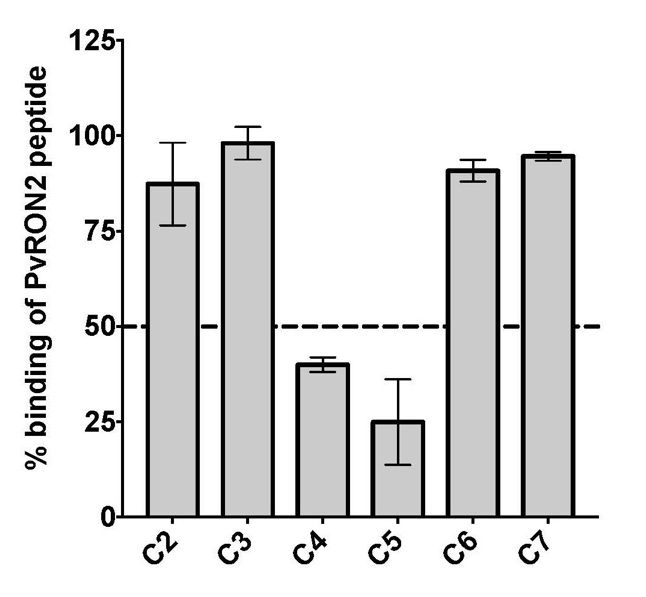

**Supplemental Figure 2. Flow diagram depicting PvAMA1-specific B cells that were isolated**.

B cells were enriched using immunomagnetic positive selection with anti-CD19 magnetic MACS beads (Miltenyi Biotec, upper left panel). SYTOX Green Dead Cell Stain (Invitrogen) was used to gate out dead cells (lower left panel). Doublet discrimination was performed to exclude aggregated cells (upper middle panel) and stained with mouse anti-human CD20 (PE-Cy5.5; Invitrogen) and anti-human IgG Abs (PE-Cy7 clone G18-145; Becton Dickinson) to identify IgG expressing B cell (lower middle panel). Pv AMA1-specific B cells were identified using biotinylated PvAMA1 using Streptavidin coupled with Fluorescein isothiocyanate (FITC) or Brilliant Violet 421 (right panel showing sort gate). PvAMA1-cells were sorted on a BD FACSAria II.

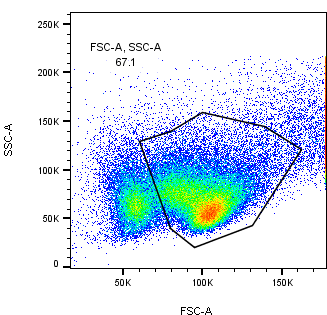

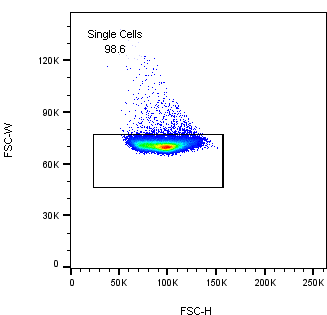

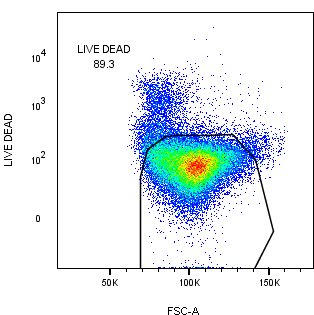

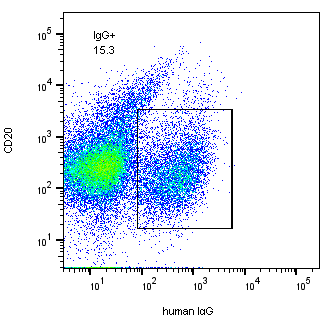

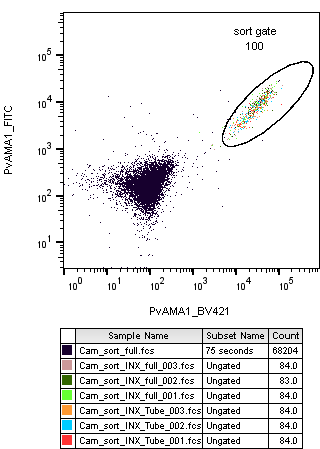

**Supplement Figure 3 – HumAb Single Cycle Kinetics Curves and Results**

A) Binding response curves to various concentrations of PvAMA1 for each humAb. **B)** k_on_ and k_off_ rates for each humAb determined using SPR.

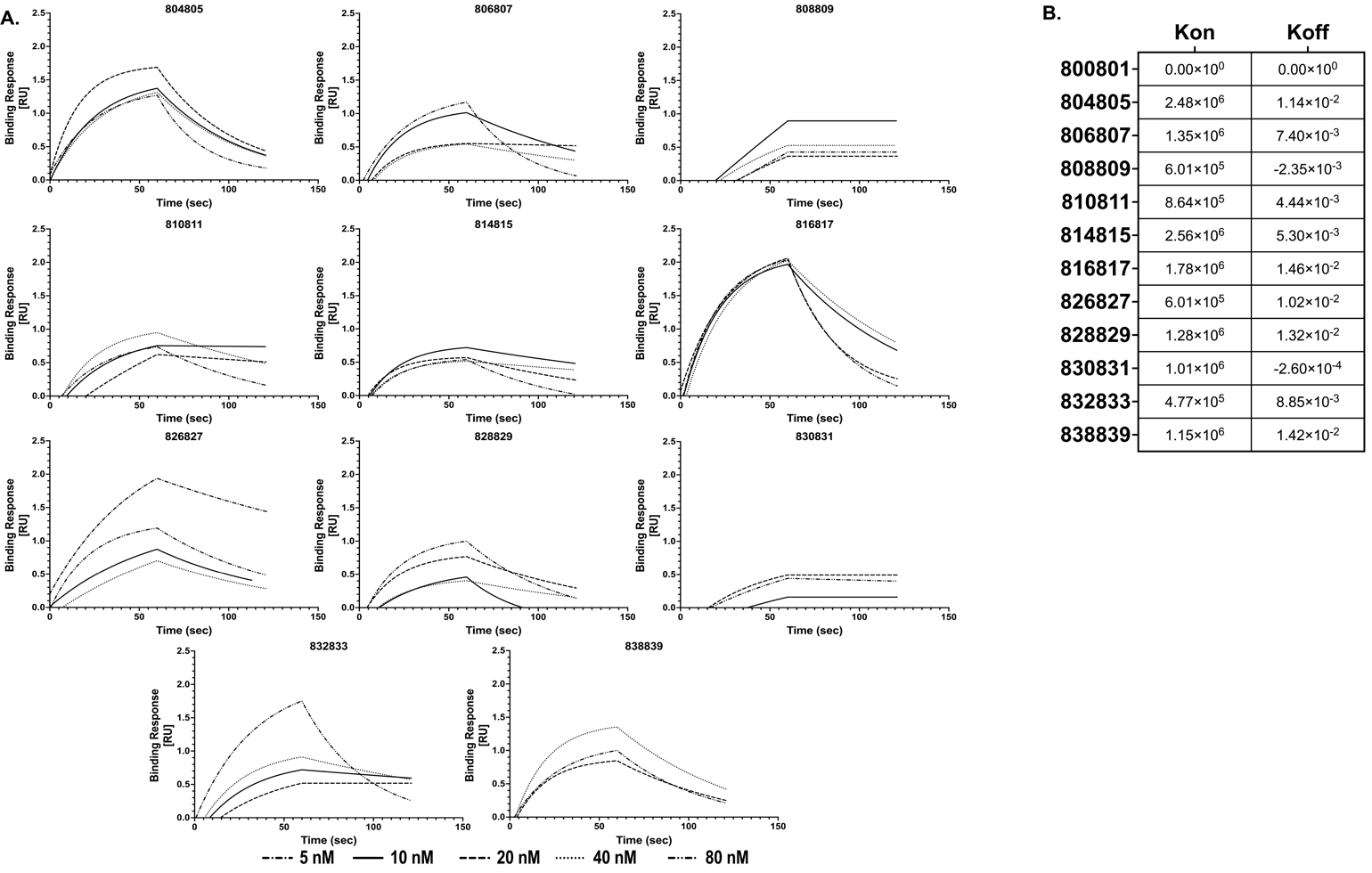

**Supplemental Figure 4 - Replicates of Pf-PvAMA1 cell line inhibition**

Dose response inhibition curves of PvAMA1-specific humAbs against Pf-PvAMA1 transgenic parasites. Each humAb was tested in triplicate as indicated in different colored lines, and IC_50_ was calculated using R. 043038, an anti-tetanus toxoid humAb was used as a negative control.

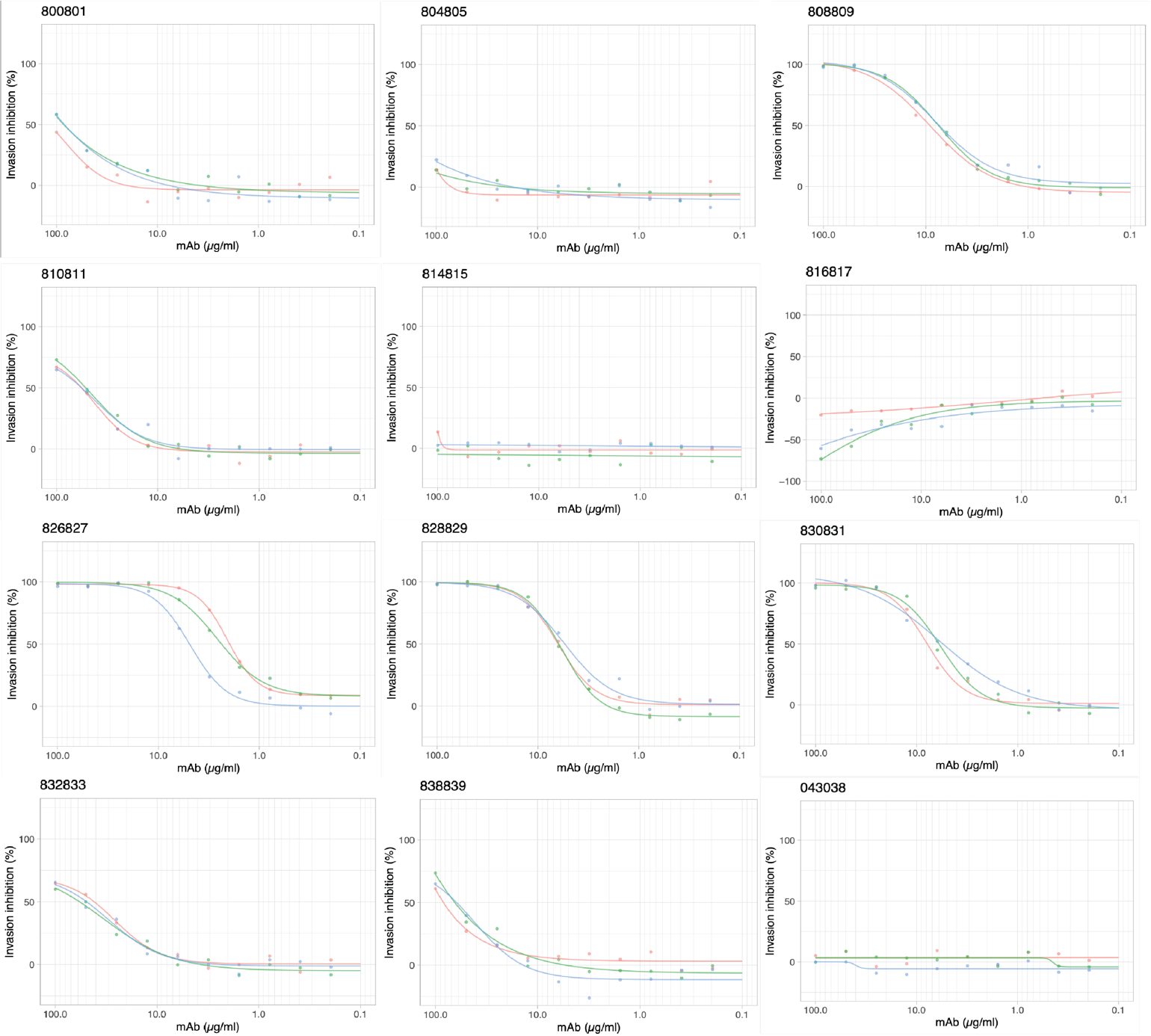

**Supplemental Figure 5 – Sporozoite HC04 invasion separated CSP210/247**

Dose response inhibition curves of Pv sporozoites blocking by PvAMA1 specific humAbs.

Murine anti-CSP monoclonal antibodies served as positive control of blocking inhibition. Three

different sporozoite isolates were used for this assay. Based on the blocking activity with the

anti-CSP monoclonal one can separate two CSP210 and one CSP247 experiments. Shown

below in **A)** are the dose-responses obtained with CSP210 strains and in **B)** with

strain CSP247. The calculated IC 50 for CSP210 is 0.08 µg/mL and for CSP247 IC 50 ~8 µg/mL

HumAbs against PvAMA1 were randomly screened with these isolates. Of note, humAb8 26827

was screened with both sporozoite strains and shows potent inhibition in both in contrast to the

CSP monoclonal.

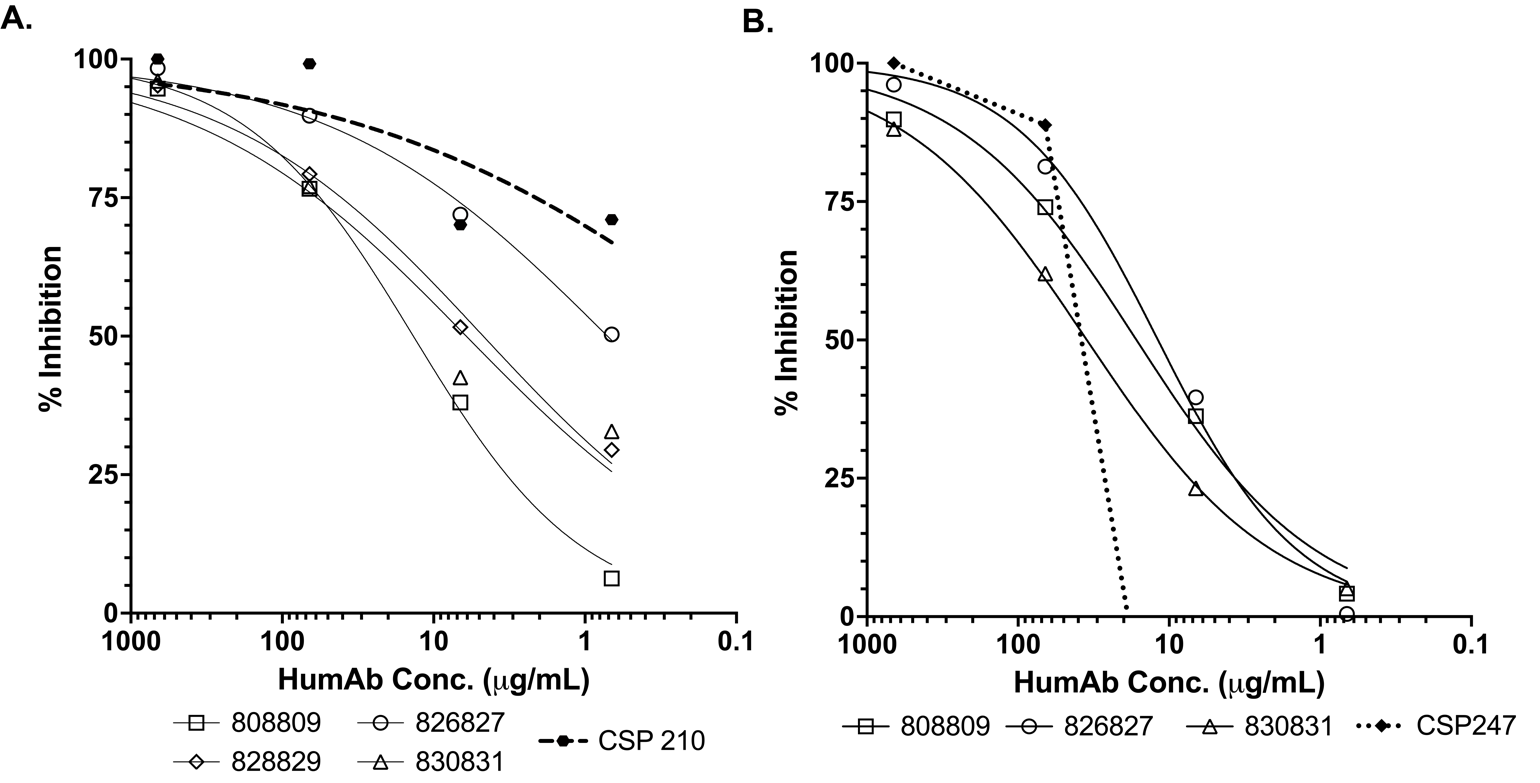

**Supplemental Figure 6 – *Microscopic assessment of P. vivax liver in FRGN huHep mice after administration of anti-AMA1 human monoclonal antibody 826827*.** The experimental design was identical to that described in the legend of Figure 4. On day 9, liver sections were analyzed microscopically for the presence of parasites described by (Mikolajczak et al. CHM, 2015). Both hypnozoites and schizonts were observed. This analysis shows total parasite forms in the liver (hypnozoites plus schizonts). Of note, one of the control livers could not be adequately evaluated and was not included in the analysis. Each dot represents one mouse. Shown in mean ± SD. Statistics: unpaired t-test.

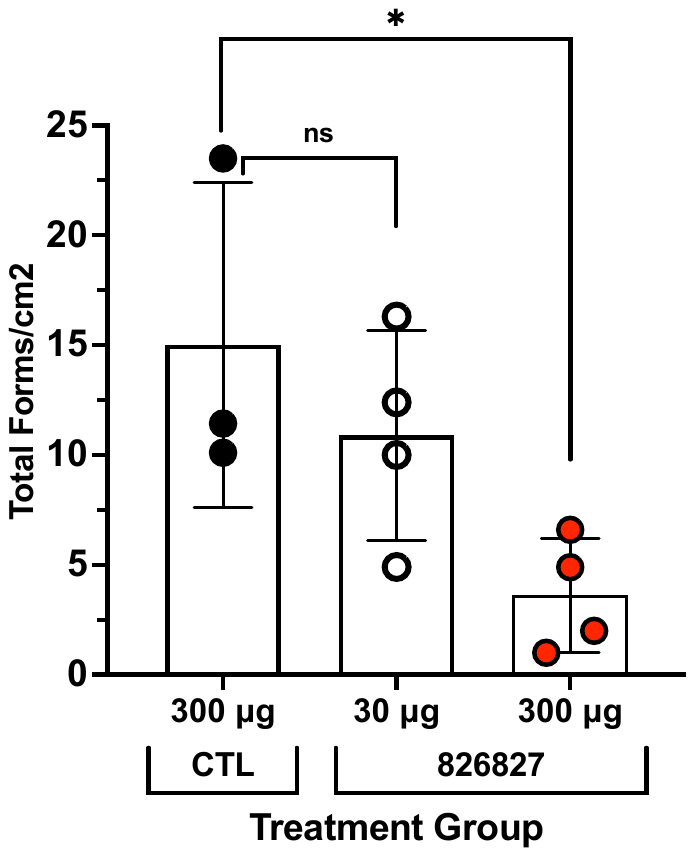

**Supplemental Figure 7 - Surface properties of PvAMA1 and its interaction partners RON2 and humAb 826827**

**A)** Schematic overview indicating the Domain 1 of PvAMA1 in gray with the Domain 2 loop in golden and the bound CDR3 loop of 826 in red. **B)** Hydrophobic surface potential of the PvAMA1 binding site. 826 was removed and rotated by 180˚ compared to the orientation in A to show the corresponding bottom interface of the interaction as well as the corresponding PvRON2 peptide. Darker areas represent higher hydrophobicity. Blue areas show hydrophilicity. **C)** Surface potential of the binding site. Red indicates negatively charged areas. Blue indicates positively charged areas. Figures were generated with Vida 4.4 from OpenEye.

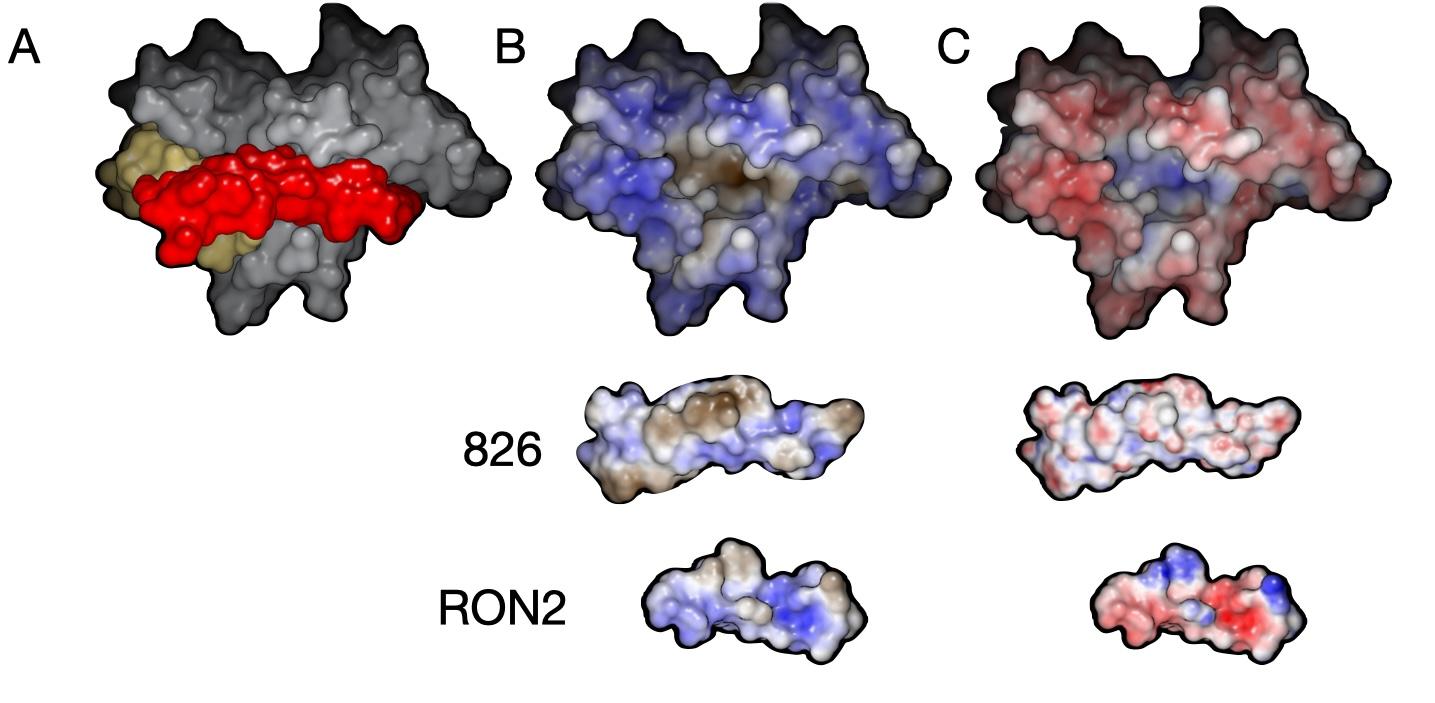

**Supplemental Figure 8 – RON2 binding groove conservation and Sequence differences between Pv, PvPNG16, Pf, Pc, Pk, Tg**

For all images, a residue-colored orange indicates an amino acid change between PvAMA1_PaloAlto (PDB: 8U9D) and another species’ version of AMA1 **A)** Sequence conservation of residues surrounding the RON2 binding groove across 390 published clinical isolates **B)** PvAMA1_PaloAlto versus PvAMA1_PNG16 **C)** PvAMA1_PaloAlto versus *P. cynomolgi* **D)** PvAMA1_PaloAlto versus *P. knowlesi* **E)** PvAMA1_PaloAlto versus *P. falciparum* **F)** PvAMA1_PaloAlto versus *Toxoplasma gondii*.

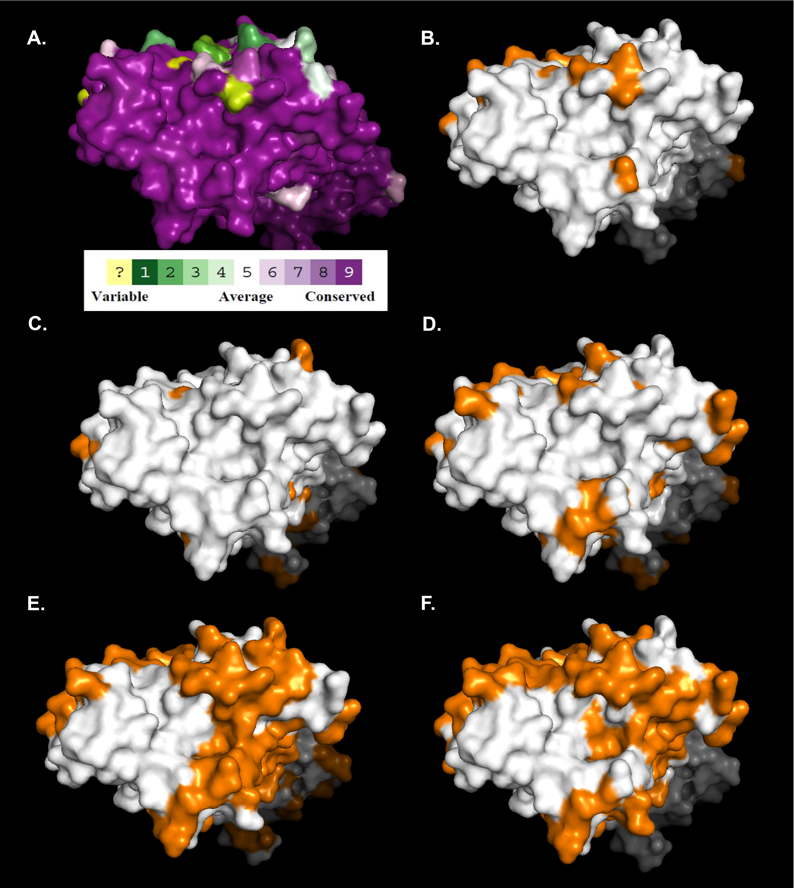

**Supplemental Figure 9 – Published PvAMA1 clinical isolates sequence conservation**

A) Percent conservation of each amino acid of PvAMA1 across 390 published clinical isolates. Yellow circles represent Domain 1 residues. Orange triangles represent Domain 2 residues. Blue squares represent Domain 3 residues. **B)** Percent conservation of PvAMA1 residues that contact the RON2 extracellular loop.

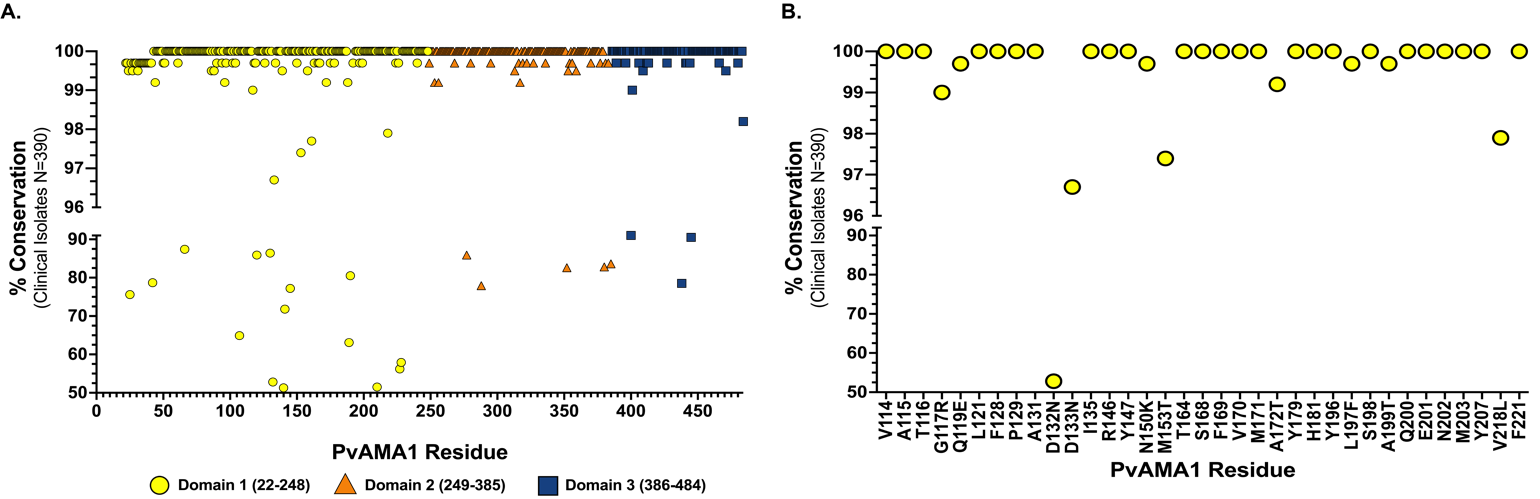

**Supplementary Figure 10. Epitope and paratope of the PvAMA1-826827 complex.**

(A) Surface representation of PvAMA1 with the epitope recognized by 826827 colored in yellow (LC contribution) and orange (HC contribution). (B) Surface representation of humAb 826827 with the paratope of PvAMA1 colored light blue (domain 1 contribution) and blue (domain 2 loop contribution). Residues within a distance cut-off of 5 Å are highlighted.

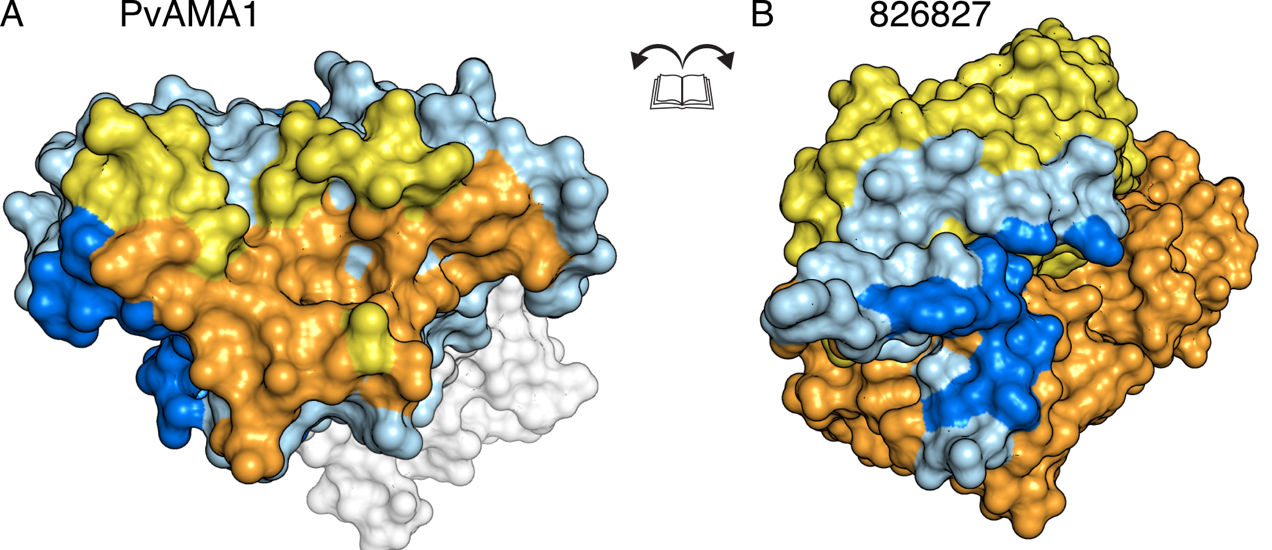

**Supplemental Table 1 – VDJ alleles and CDR3 sequences and clonal groups**

Shown are VDJ alleles and CDR3 sequences for 157 sorted and sequenced Bc IgH. The first column shows B cells that have been made into monoclonal antibodies, assigned a number, and the number of B cells in the clonal group.

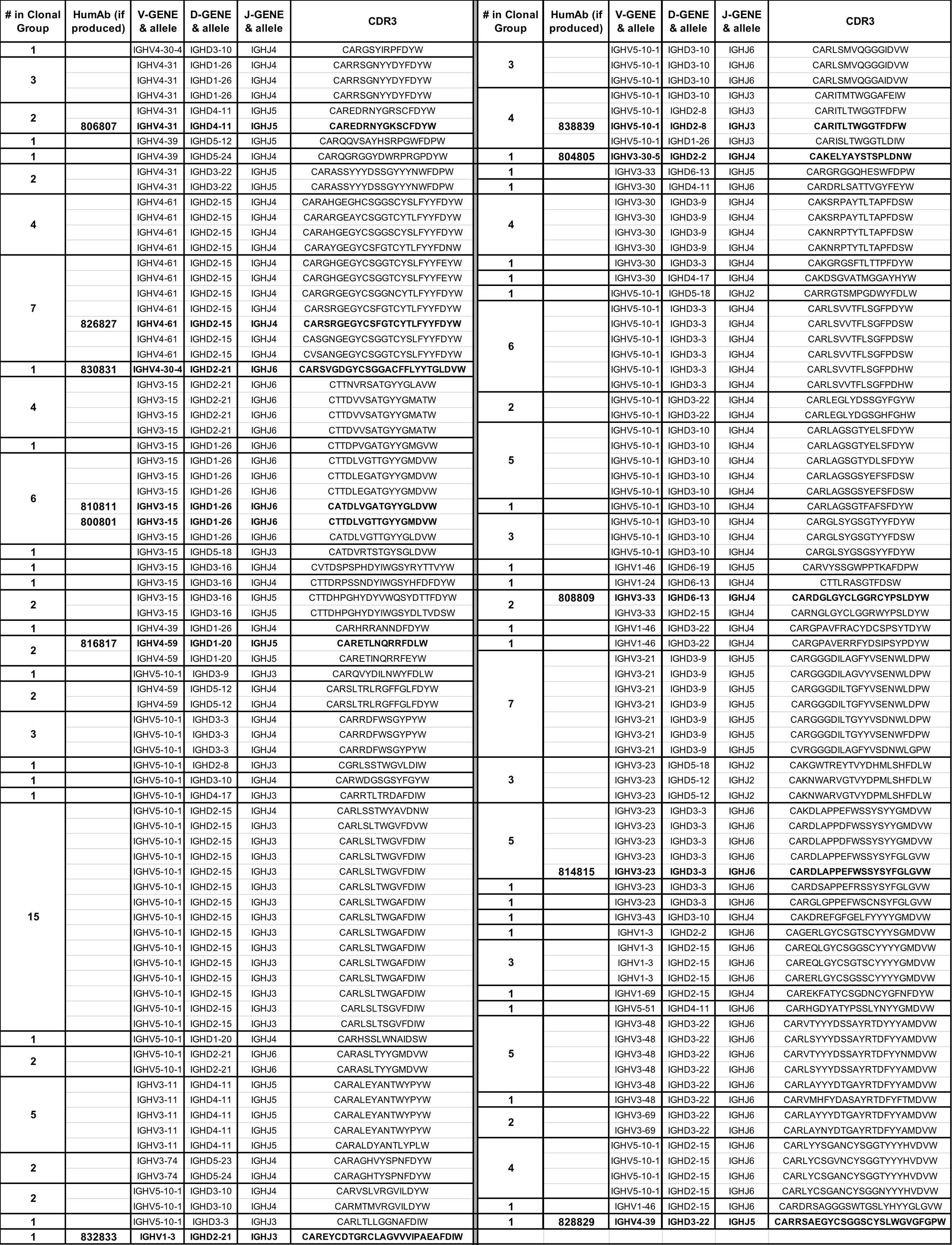

**Supplemental Table 2 – Avidity Index (AI_50_) of PvAMA1 specific humAbs to different AMA1 constructs**

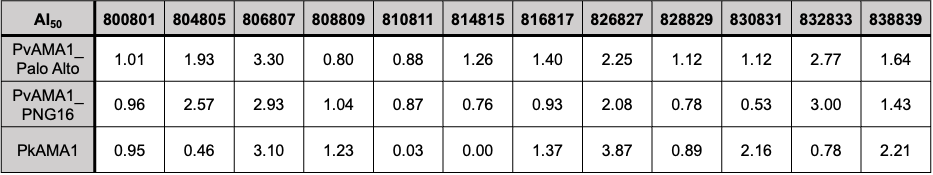

**Supplemental Table 3 - List of SNP and haplotypes observed in AMA1 sequences in isolates tested in response to 826827**

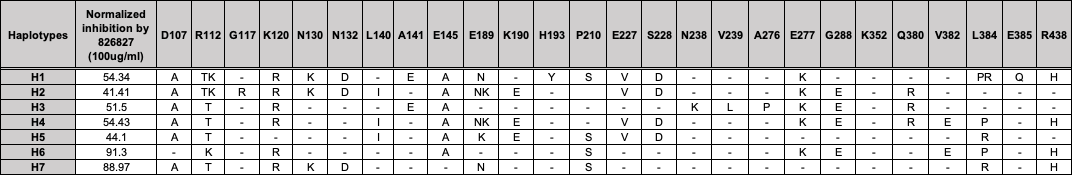

**Supplemental Table 4 – Crystallography data collection and refinement statistics**

|  | PvAMA1-Fab 826827 |
| --- | --- |
| PDB ID | 8U9D |
| **Data collection statistics** |  |
| Wavelength (Å) | 0.953725 |
| Space group | C121 |
| Cell axes (Å) (a, b, c) | 187.24, 54.46, 104.13 |
| Cell angles (º) (α, γ, β) | 90, 97.20, 90 |
| Resolution range (Å) | 47.76-2.40  (2.54-2.40)* |
| Completeness (%) | 99.9 (99.8) |
| Total no. of reflections | 289692 (45358) |
| Unique reflections | 41378 (6614) |
| Redundancy | 7.0 (6.9) |
| R_meas_ (%) | 18.0 (117.6) |
| CC_1/2_ (%) | 99.5 (64.2) |
| I/σ | 9.53 (1.55) |
| Wilson B (Å^2^) | 42.79 |
| **Refinement statistics** |  |
| R_work_/R_free_ (%) | 19.2/ 23.5 |
| No. of atoms |  |
| Protein | 6395 |
| Water | 259 |
| B factors (Å^2^) |  |
| Chain A | 46.8 |
| Chain B | 45.3 |
| Chain C | 49.6 |
| Water | 45.7 |
| R.m.s. deviations |  |
| Bond lengths (Å) | 0.004 |
| Bond angles (º) | 0.668 |
| Validation |  |
| Ramachandran plot |  |
| outliers (%) | 0.0 |
| favored (%) | 96.6 |
| Rotamer outliers (%) | 0.9 |
| C-beta outliers | 0 |
| MolProbity score | 1.49 |

* The values in parentheses represent the highest-resolution shell.

**Supplemental Table 5 - Interactions PvAMA1 – Fab 826827 based on PISA**

| PvAMA1 | Group | 826827 | Location | Group | Distance |
| --- | --- | --- | --- | --- | --- |
| Hydrogen bonds | | | | | |
| Glu 83 | OE2 | Tyr 54 | CDR-H2 | OH | 2.4 |
| Glu 83 | OE2 | Tyr 55 | CDR-H2 | OH | 3.3 |
| Thr 116 | OG1 | Tyr 49 | CDR-L2 | OH | 2.9 |
| Asp 118 | OD1 | Gly 57 | CDR-L2 | N | 3.0 |
| Asp 118 | OD2 | Arg 54 | CDR-L2 | NH2 | 2.8 |
| Asp 118 | N | Arg 54 | CDR-L2 | O | 2.9 |
| Asp 118 | O | Arg 54 | CDR-L2 | NH2 | 3.7 |
| Ala 131 | O | Lys 31 | CDR-L1 | N | 2.9 |
| Ala 131 | O | Thr 32 | CDR-L1 | N | 3.5 |
| Asn 132 | ND2 | Thr 113 | CDR-H3 | O/N | 2.8/ 3.5 |
| Asp 133 | N | Ser 30 | CDR-L1 | OG | 2.9 |
| Val 170 | O | Cys 106 | CDR-H3 | N | 2.7 |
| Val 170 | N | Cys 106 | CDR-H3 | O | 2.7 |
| Ala 172 | N | Gly 104 | CDR-H3 | O | 2.9 |
| Tyr 196 | OH | Arg 101 | CDR-H3 | NH2 | 3.5 |
| Tyr 196 | OH | Glu 103 | CDR-H3 | OE1/OE2 | 2.8/2.6 |
| Gln 314 | O | Tyr 34 | CDR-H1 | OH | 3.6 |
| Asn 315 | ND2 | Tyr 117 | CDR-H3 | OH | 2.8 |
| Asn 315 | O | Arg 101 | CDR-H3 | NE | 2.7 |
| Asn 315 | O | Tyr34 | CDR-H1 | OH | 3.8 |
| Asn 315 | OD1 | Arg 101 | CDR-H3 | NH2 | 3.6 |
| Asn 316 | ND2 | Tyr 34 | CDR-H1 | OH | 3.7 |
| Lys 321 | NZ | Tyr 105 | CDR-H3 | OH | 2.6 |
| Salt bridges | | | | | |
| Lys 321 | NZ | Glu 103 | CDR-H3 | OE2 | 3.6 |
| Asp 118 | OD2 | Arg 54 | CDR-L2 | NH2 | 2.8 |
| Other interfacing residues in PvAMA1 | | | | | |
| Asn 84 | Gly 117 | Gln 119 | Phe 128 | Pro 129 | Asn 130 |
| His 134 | Ile 135 | Arg 146 | Tyr 147 | Asn 150 | Met 153 |
| Thr 164 | His 165 | Ser 168 | Phe 169 | Met 171 | Gly 173 |
| Gln 175 | His 181 | Met 194 | Gln 310 | Arg 313 | Arg 317 |
| Glu 318 |  |  |  |  |  |
| Other interfacing residues in 826 (heavy chain) | | | Other interfacing residues in 827 (light chain) | | |
| Ser 331 | Pro 32 | Gly 33 | Ser 28 | Ser 50 | Thr 53 |
| Tyr 35 | Arg 56 | Arg 99 | Ala 55 | Ser 56 | Val 58 |
| Gly 102 | Ser 107 | Phe 108 | Tyr 91 | Asn 92 | Trp 94 |
| Cys 111 | Tyr 112 | Leu 114 |  |  |  |
| Phe 115 | Asp 119 | Tyr 120 |  |  |  |

**Supplemental Movie 1: PvAMA1 bound Fab826827 compared to RON2**

The electrostatic surface potential of PvAMA1 (8U9D) is displayed without 826827 bound to it, then the superimposed RON2 is shown in magenta followed by the 826 CD3 loop in green, showing a tight overlap in the binding site. Next the backbone of 826 and 827 are displayed to show our complete co-crystal structure.

**Supplemental Movie 2: Polymorphisms within PvAMA1 in relation to Fab826827** The co-crystals structure of PvAMA1 with humAb 826827 is shown as an overview where PvAMA1 is represented as a solid surface and 826 in blue and 827 in yellow. The movie zooms towards three residues of interest namely D132, N130 and G117 which represent the wildtype amino acid residues and then these residues switch to the mutations observed in our clinical isolates as well as in the known sequences from other clinical isolates.

**Supplemental Movie 3: Morph between open and closed conformation of AMA1 Domain 2 loop**

A morph between PDB ID 6N87 and 8U9D is shown that focuses on the Domain 2 loop movement. The PfAMA1 structure 6N87 was superimposed onto 8U9D prior to generating the intermediate states for the moving Domain 2 loop.
